# *In vitro* evaluation of the activity of terpenes and cannabidiol against Human Coronavirus E229

**DOI:** 10.1101/2021.03.01.433501

**Authors:** Lior Chatow, Adi Nudel, Iris Nesher, David Hayo Hemo, Perri Rozenberg, Hanna Voropaev, Ilan Winkler, Ronnie Levy, Zohar Kerem, Zohara Yaniv, Nadav Eyal

## Abstract

The activity of a new, terpene-based formulation, code-named NT-VRL-1, against Human Coronavirus (HCoV) strain 229E was evaluated in human lung fibroblasts (MRC-5 cells), with and without the addition of cannabidiol (CBD). The tested formulation exhibited an antiviral effect when it was pre-incubated with the host cells prior to virus infection. The combination of NT-VRL-1 with CBD potentiated the antiviral effect better than the positive controls pyrazofurin and glycyrrhizin. There was a strong correlation between the quantitative results from a cell-viability assay and the cytopathic effect seen under the microscope after 72 h. To the best of our knowledge, this is the first report of activity of a combination of terpenes and CBD against a coronavirus.

## Introduction

Coronaviruses are enveloped, non-segmented, positive-strand RNA viruses of the family Coronaviridae that cause a wide spectrum of illnesses in humans, including respiratory and gastrointestinal diseases (1). To date, seven human coronaviruses (HCoVs) have been identified. Four of those, HCoV-229E, HCoV-OC43, HCoV-NL63 and HCoV-HKU1, are non-zoonotic and cause worldwide outbreaks of upper respiratory tract infections predominantly in the winter (2). The severe acute respiratory syndrome coronavirus 2 (SARS-CoV-2) has produced an epidemic of Coronavirus Disease 2019 (COVID-19) that started on 31 December 2019 in China and then spread to different regions and countries. According to the WHO (3), as of January 26th, 2021, there have been a total of 98,925,221 confirmed cases of COVID-19 worldwide, including 2,127,294 deaths.

This outbreak has led to a search for active antiviral compounds to treat this disease. While SARS-CoV-2 is highly contagious and can only be studied in a biosafety level 4 facility, working with the less virulent strain HCoV-229E is considered a good alternative for preliminary research (2, 4, 5). HCoV-229E is associated with various respiratory illnesses ranging from the common cold to severe pneumonia (6). Recently, the potential of phytochemicals, such as terpenes, for use as potent antiviral agents has received considerable attention, especially because these substances are naturally abundant with relatively low toxicity and cost (7).

Terpenes are natural, volatile compounds primarily extracted from plants, which contain only carbon, hydrogen and oxygen atoms. In plants, terpenes act as chemoattractants or chemorepellents (8) and are largely responsible for plant fragrances. In animals and humans, terpenes exhibit a variety of pharmacological properties, including anti-inflammatory (9), analgesic (10), antimicrobial (11) and antiviral (12) properties. A wide range of *in vitro* studies have demonstrated terpenes’ potential for use against a wide range of viruses such as herpes simplex virus (13), bronchitis virus (14), West Nile virus (15) and HIV-1 (16).

As isolated compounds and in plant essential oils, terpenes have been shown to have antiviral effects against several types of HCoVs. Glycyrrhizin, a triterpene found in licorice roots, was one of the first compounds found to be active against SARS coronavirus (SARS-CoV) *in vitro*; it was shown to inhibit SARS-CoV replication with an EC_50_ of 365 μM (17). Glycyrrhizin has also been used to successfully treat SARS patients (18). *Laurus nobilis* essential oil, with beta-ocimene, 1,8-cineole, alpha-pinene and beta-pinene as its main constituents, was found to exert antiviral activity against SARS-CoV with an IC_50_ value of 120 mg/mL (19).

Even though the vaccination of the world’s population against COVID-19 has begun and is expected to proceed gradually, with no clear expectation of completion. Some individuals will not be vaccinated due to personal choice or health limitations. In addition, several population groups such as younger age groups will be the last to get vaccinated. A natural antiviral solution with minimal side effects that can be used alone or in conjunction with vaccines as a preventative treatment may be a safe and relatively easy way to reduce infection in those populations.

The goal of the present study was to evaluate the antiviral activity of a proprietary terpene formulation (code named NT-VRL-1) against HCoV-229E, with and without the addition of cannabidiol (CBD), and the mode of antiviral action of this formulation during the viral multiplication cycle. The NT-VRL-1 formulation consisted of 30 natural terpenes that are found in cannabis, as well as other plants. The therapeutic activity of these compounds was evaluated in terms of the cytopathic effect observed under an inverted microscope and an *in vitro* cell viability XTT assay involving human lung fibroblasts (MRC-5 cells) in which the mitochondrial activity of those cells was examined.

## Materials and Methods

### Materials and reagents

MRC-5 cells and the HCoV-229E strain were purchased from the American Type Culture Collection (ATCC; Manassas, Virginia, United States). All media ingredients and the XTT-based viability assay kit were purchased from Biological Industries (Beit HaEmek, Israel). CBD was purchased from Recipharm Israel (Ness Ziona, Israel). NT-VRL-1 was obtained from Eybna Technologies (Givat Hen, Israel). Glycyrrhizin was obtained from Penta International Corporation (New Jersey, USA) and pyrazofurin was purchased from Sigma (Jerusalem, Israel).

### Cytotoxicity of compounds

MRC-5 cells were plated at 1 × 104 cells/well in 96-well plates in minimum essential medium Eagle (EMEM) supplemented with 10% fetal calf serum and then incubated at 37°C with 5% CO2. The next day, the medium was discarded and 100 μL of EMEM supplemented with 1% fetal calf serum was added to the cells, together with the compounds. The following concentrations were tested for each potential treatment. CBD: 2 μg/mL, 5 μg/mL and 10 μg/mL. NT-VRL-1: 5 μg/mL, 10 μg/mL, 50 μg/mL and 100 μg/mL. NT-VRL-1 + CBD: 10 μg/mL + 1 μg/mL and 10 μg/mL + 3 μg/mL (respectively). Pyrazofurin: 2 μg/mL, 5 μg/mL and 10 μg/mL. Glycerrihizin: 100 μg/mL, 500 μg/mL and 1000 μg/mL. The cells were incubated for an additional 72 ± 2 h at 34°C and 5% CO2. Finally, the cells were subjected to an XTT assay. Based on the results of this work, we determined the non-toxic concentrations of the compounds to be used in the efficacy evaluations: CBD (0.5 μg/mL and 1 μg/mL), NT-VRL-1 (2 μg/mL, 5 μg/mL and 10 μg/mL), NT-VRL-1 + CBD (10 μg/mL + 1 μg/mL), pyrazofurin (5 μg/mL) and glycerrihizin (400 μg/mL).

### Efficacy of compounds - Cell pretreatment

MRC-5 cells were plated at 1 × 10^4^ cells/well in 96-well plates in EMEM supplemented with 10% fetal calf serum, and then incubated at 37°C and 5% CO_2_. The next day, the medium was discarded and 100 μL of EMEM supplemented with 1% fetal calf serum was added to the cells, supplemented with the compounds at concentrations previously identified as nontoxic. The cells were incubated for 1 h at 34°C and 5% CO2. Next, 1 μL of medium or virus at 100 times the concentration of the infective dose (1:340 dilution) was added to the cells. The cells were incubated for an additional 72 ± 2 h at 34°C and 5% CO_2_. Under an inverted microscope, a photograph was taken of the cells in each treatment at 24, 48 and 72 h post-infection. A virus-induced cytopathic effect was observed in comparison with the parallel virus control and cell control. Finally, cells were subjected to an XTT assay.

### Efficacy of compounds - Virus pretreatment

MRC-5 cells were plated at 1 × 10^4^ cells/well in 96-well plates in EMEM supplemented with 10% fetal calf serum, and then incubated at 37°C and 5% CO_2_. The next day, 120 μL of EMEM supplemented with 1% fetal calf serum was added to the wells, supplemented with the compounds at concentrations previously identified as nontoxic. The virus was mixed with the compounds in a U-shaped plate and then incubated for 1 h at 34°C and 5% CO_2_. Then, 1.2 μL medium or virus at 100 times the concentration of the infective dose was added to the wells. Next, 100 μL of the virus + compounds mixture was added to the cells after medium was removed and the cells were incubated at 34°C and 5% CO_2_ for an additional 72 ± 2 h. Under an inverted microscope, photographs of the cells in each treatment were taken at 24, 48 and 72 h post-infection. The virus-induced cytopathic effect was observed in comparison with the parallel virus control and cell control. Finally, cells were subjected to an XTT assay.

### Efficacy of compounds – Post-adsorption

MRC-5 cells were plated at 1 × 10^4^ cells/well in 96-well plates in EMEM supplemented with 10% fetal calf serum, and then incubated at 37°C and 5% CO_2_. The next day, medium was discarded and EMEM supplemented with 1% fetal calf serum was added to the cells with or without 1 μL of virus at 100 times the concentration of the infective dose. The cells were then incubated for 1 h at 34°C and 5% CO_2_. Then, medium was discarded and 100 μL of EMEM supplemented with 1% fetal calf serum was added to the cells, supplemented with the compounds at the concentrations previously identified as nontoxic and 1 μL of media or virus at 100 times the concentration of the infective dose. The cells were incubated for an additional 72 ± 2 h at 34°C and 5% CO_2_. Under an inverted microscope, photographs were taken of the cells in each treatment at 24, 48 and 72 h post-infection. The virus-induced cytopathic effect was observed in comparison with the parallel virus control and cell control. Finally, cells were subjected to an XTT assay.

### XTT-based viability assay

At the end of each incubation period, media was discarded from all wells and 100 μL of fresh culture medium was added to the cells together with 50 μL of XTT reagent. OD was measured at 450 nm (after subtraction of the non-specific OD at 620 nm).

## Results

### Cytotoxicity of compounds

The non-cytotoxic concentrations of the various compounds were determined as the concentrations that did not lead to excess cell death, as compared to untreated cells. As shown in **Figure 1**, the non-toxic concentrations were: CBD ≤ 1 μg/mL, NT-VRL-1 ≤ 10 μg/mL, pyrazofurin ≤ 10 μg/mL and glycyrrhizin ≤ 500 μg/mL.

**Figure 1:**
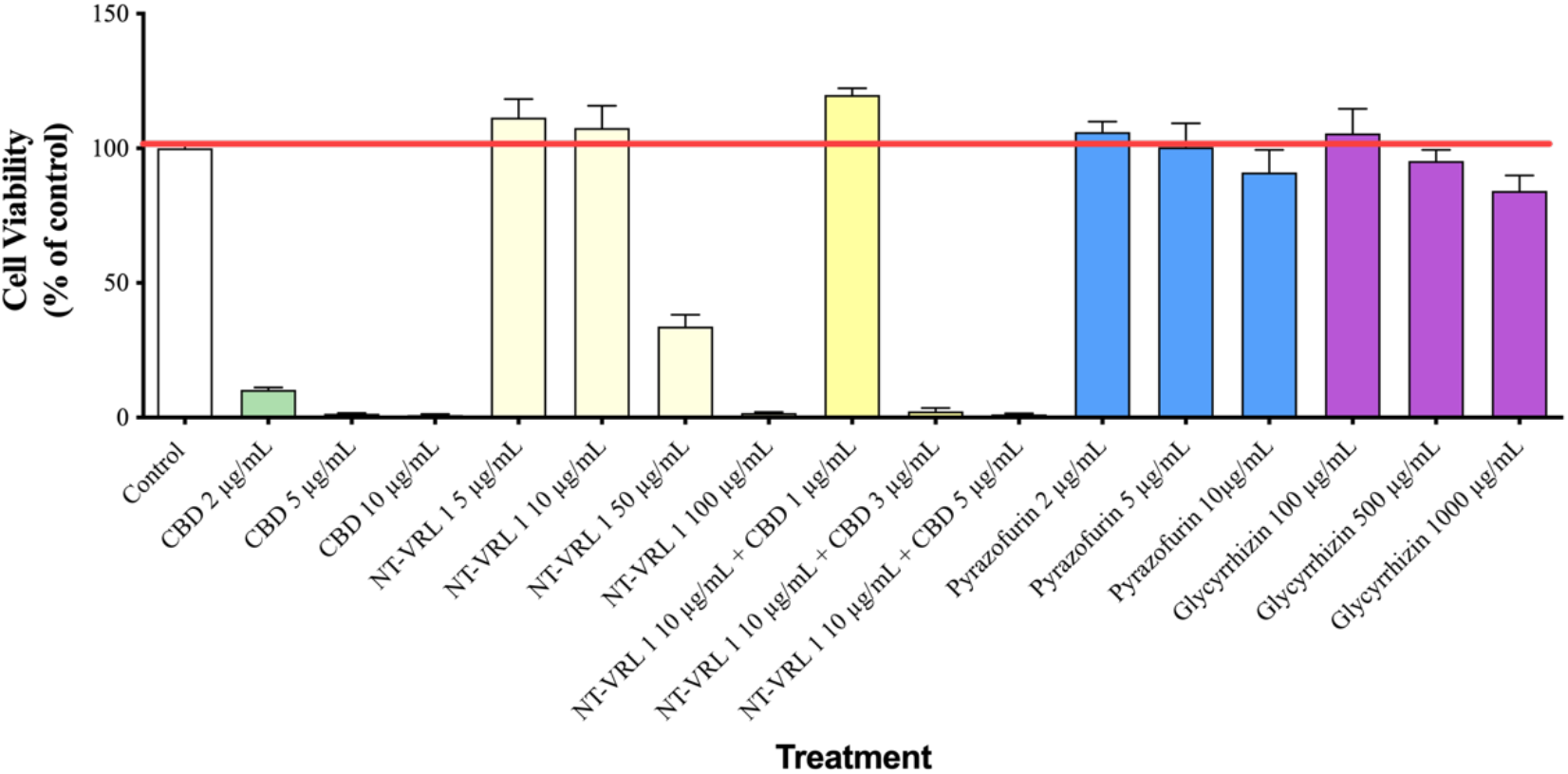
Cytotoxicity test. MRC-5 cells were treated with different concentrations of the compounds for 72 h. Cell viability was then determined using an XTT assay. Results represent mean percent viability ± SEM (*n* = 4).

### Efficacy of compounds - Cell pretreatment

MRC-5 cells were pretreated with the compounds prior to inoculation with HCoV-229E. As shown in **Figure 2**, the viability of cells that were infected with HCoV-229E, but otherwise untreated, was reduced to ~40% of the viability of the uninfected control cells. Pre-incubation of the cells with all of the compounds prior to virus inoculation rescued the cells and increased the level of cell viability. The combination of 10 μg/mL NT-VRL-1 with 1 μg/mL CBD was the most effective treatment associated with the highest level of cell viability (p < 0.001). This pattern was also observed in terms of the cytopathic effect seen under the microscope after 72 h. Swelling and clumping of the MRC-5 cells was observed 72 h after viral infection (**Figure 3**). Cell pretreatment with NT-VRL-1 alone (**Figure 3C**) or NT-VRL-1 + CBD (**Figure 3D**) before viral infection prevented a cytopathic effect.

**Figure 2:**
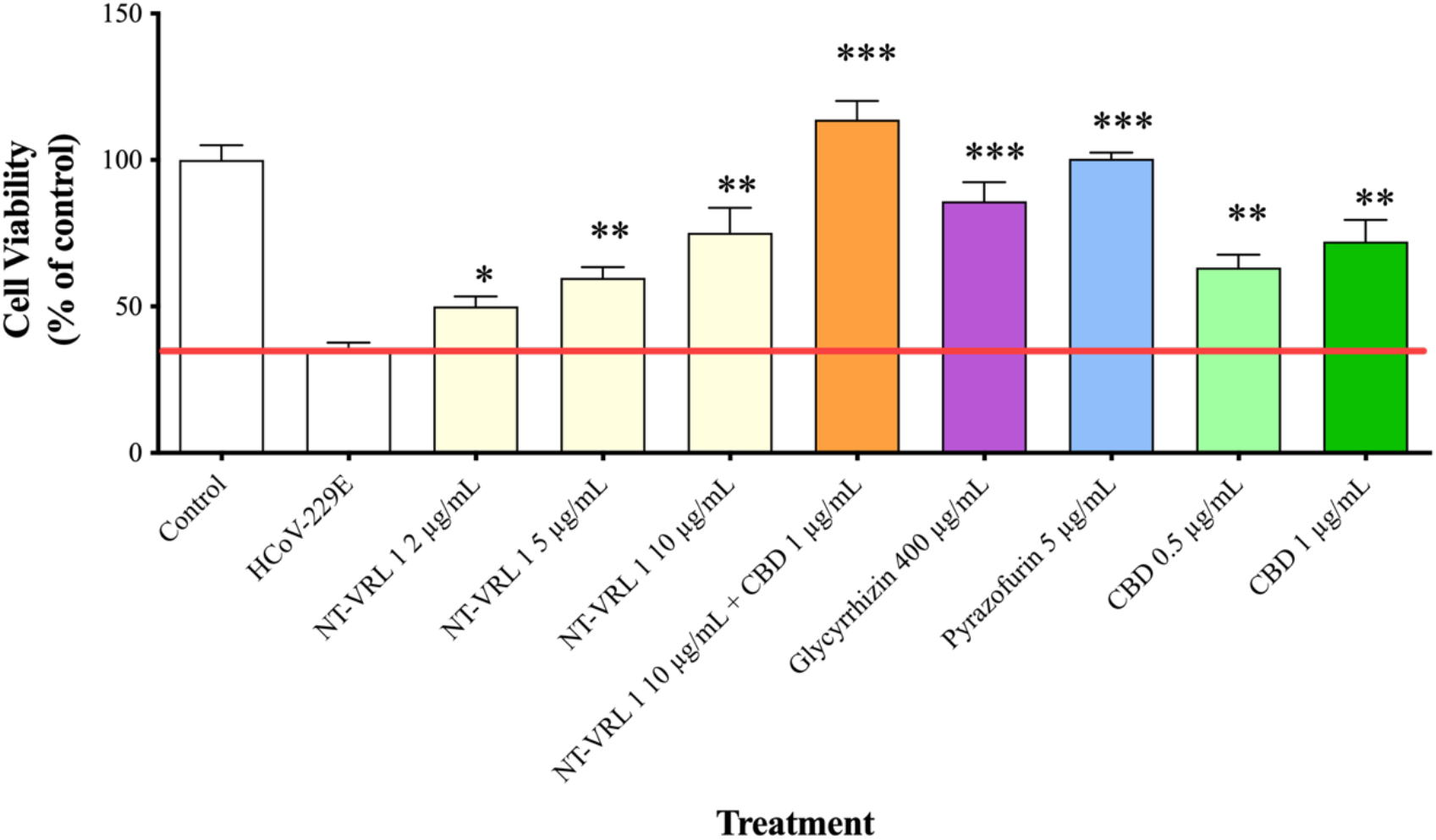
Protective effect of pretreatment of MRC-5 cells with the compounds against HCoV-229E infection. MRC-5 cells were first pretreated with different concentrations of the compounds for 1 h and then exposed to HCoV-229E for an additional 72 h. Cell viability was determined using an XTT assay. Results represent mean percent viability ± SEM (*n* = 4). Statistics are presented for each treatment compared to cells treated with HCoV-229E only. \**p* < 0.05, \*\**p* < 0.01 and \*\*\**p* < 0.001, according to a *t*-test.

**Figure 3:**
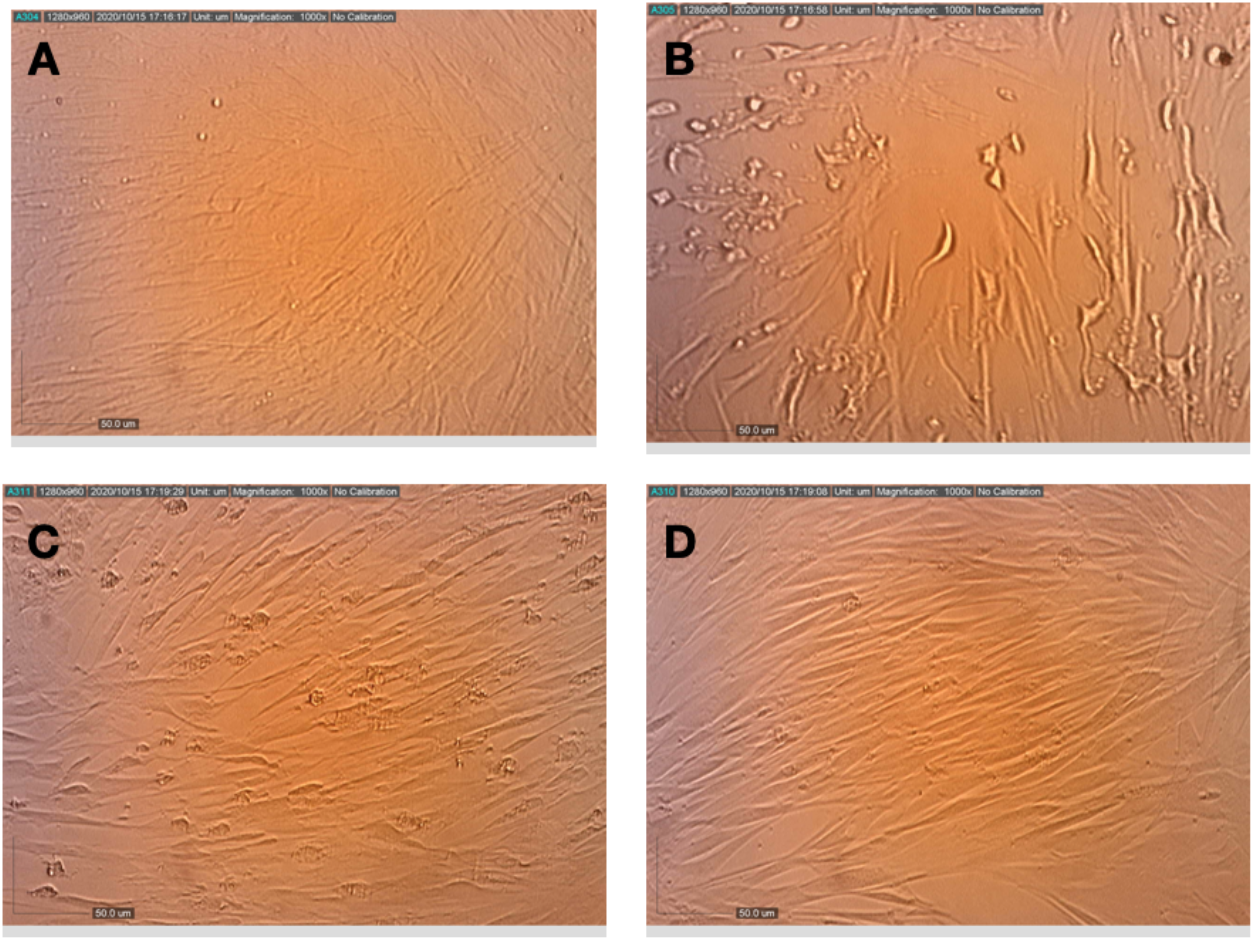
Effects of pretreatment of MRC-5 cells with terpenes and CBD on the replication and cytopathic effect of HCoV-229E. (A) Healthy MRC-5 cells, (B) MRC-5 cells that had been pretreated with assay medium, photographed 72 h after inoculation with HCoV-229E, (C) MRC-5 cells that had been pretreated with terpenes, photographed 72 h after infection with HCoV-229E, (D) MRC-5 cells that had been pretreated with terpenes and CBD, photographed at 72 h after infection with HCoV-229E.

### Efficacy of compounds - Virus pretreatment

HCoV-229E was incubated with the compounds before it was introduced to the MRC-5 cells. As shown in **Figure 4**, inoculation with HCoV-229E (pre-incubation with assay medium) reduced the viability to 80%. Pre-incubation of the virus with 10 μg/mL NT-VRL-1 + 1 μg/mL CBD prior to its introduction to host cells elevated cell viability back to the level observed for the control (p < 0.001). In addition, virus pretreatment with NT-VRL-1 + CBD prevented a cytopathic effect after the cells were inoculated with the virus (**Figure 5**)

**Figure 4:**
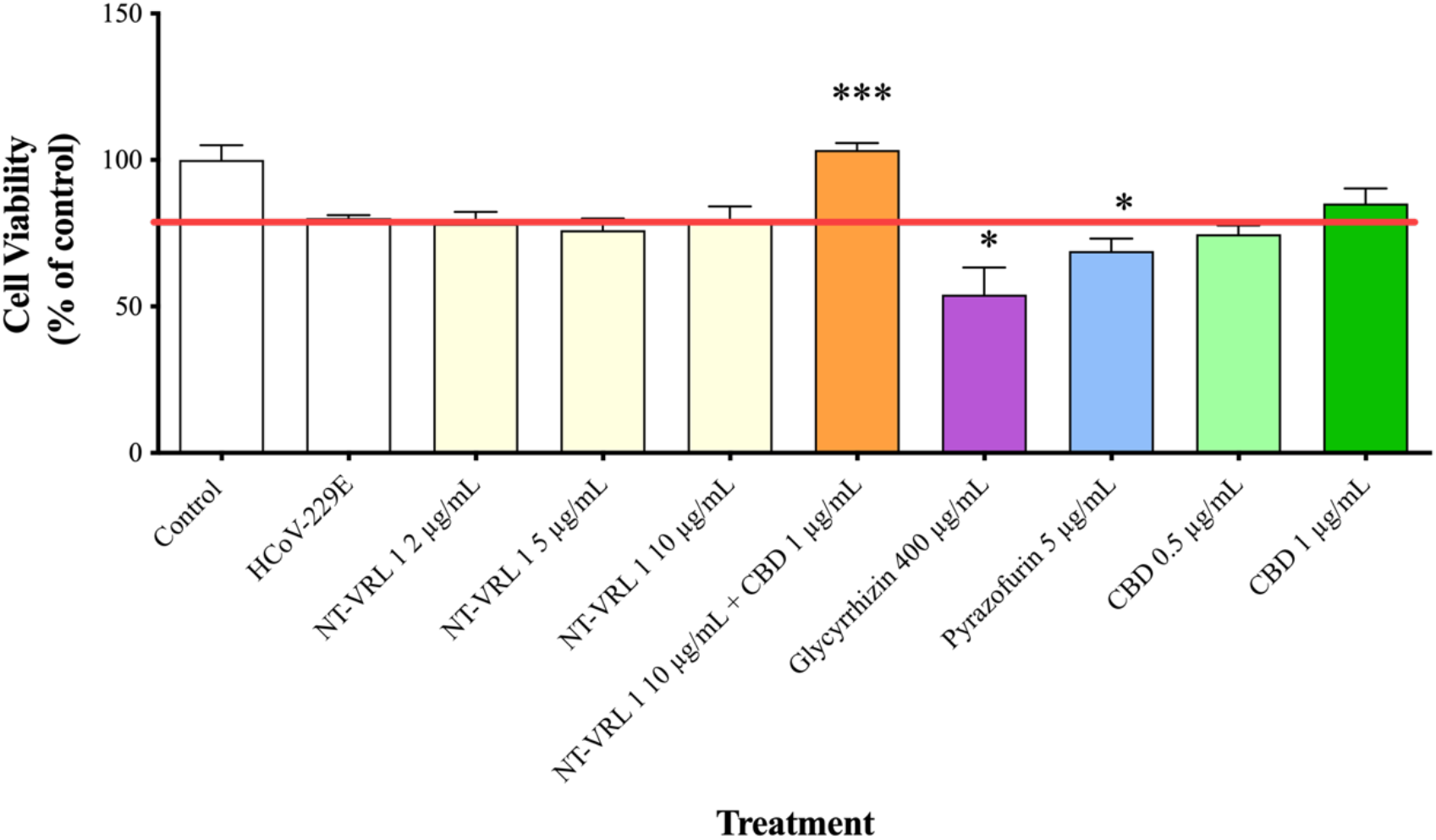
Effect of pretreatment of HCoV-229E with the compounds on the viability of MRC-5 cells. HCoV-229E was treated with different concentrations of the compounds for 1 h and then incubated with MRC-5 cells for an additional 72 h. Cell viability was determined using an XTT assay. Results represent mean percent viability ± SEM (*n* = 4). Statistics are presented for each treatment relative to cells treated with HCoV-229E that had not been pretreated. \**p* < 0.05 and \*\*\**p* < 0.001, according to a *t*-test.

**Figure 5:**
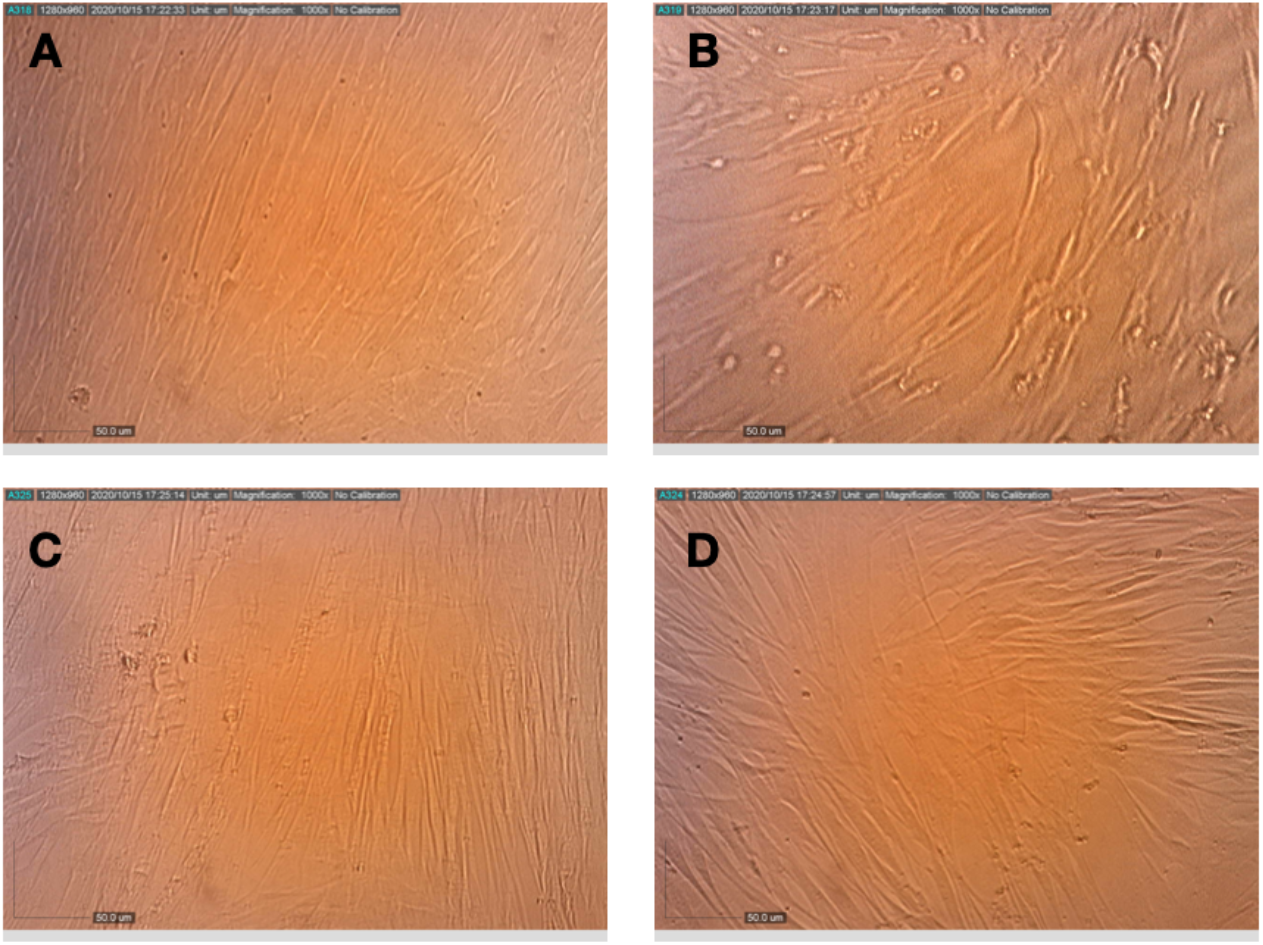
Effect of pretreatment of HCoV-229E with terpenes and CBD on its replication and cytopathic effect in MRC-5 cells. (A) Healthy MRC-5 cells, (B) MRC-5 cells at 72 h after infection with HCoV-229E that had been pretreated with assay medium, (C) MRC-5 cells at 72 h after infection with HCoV-229E that had been pretreated with terpenes and(D) MRC-5 cells at 72 h after infection with HCoV-229E that had been pretreated with terpenes and CBD.

### Efficacy of compounds – Post-adsorption

When the compounds were added to the cells after virus adsorption, the viability of the HCoV-229E-infected cells was reduced to only ~70% of the control, as shown in **Figure 6**. Under these conditions, 10 μg/mL NT-VRL-1 + 1 μg/mL CBD prevented cell death and preserved a level of cell viability similar to that observed for the control (p < 0.001). Pyrazofurin at 5 μg/mL also enhanced cell viability (p < 0.05) relative to the untreated, infected control. Similar results were observed in terms of the cytopathic effect after 72 h, at which point NT-VRL-1 by itself (**Figure 7C**) and NT-VRL-1 + CBD (**Figure 7D**) both prevented cell damage.

**Figure 6:**
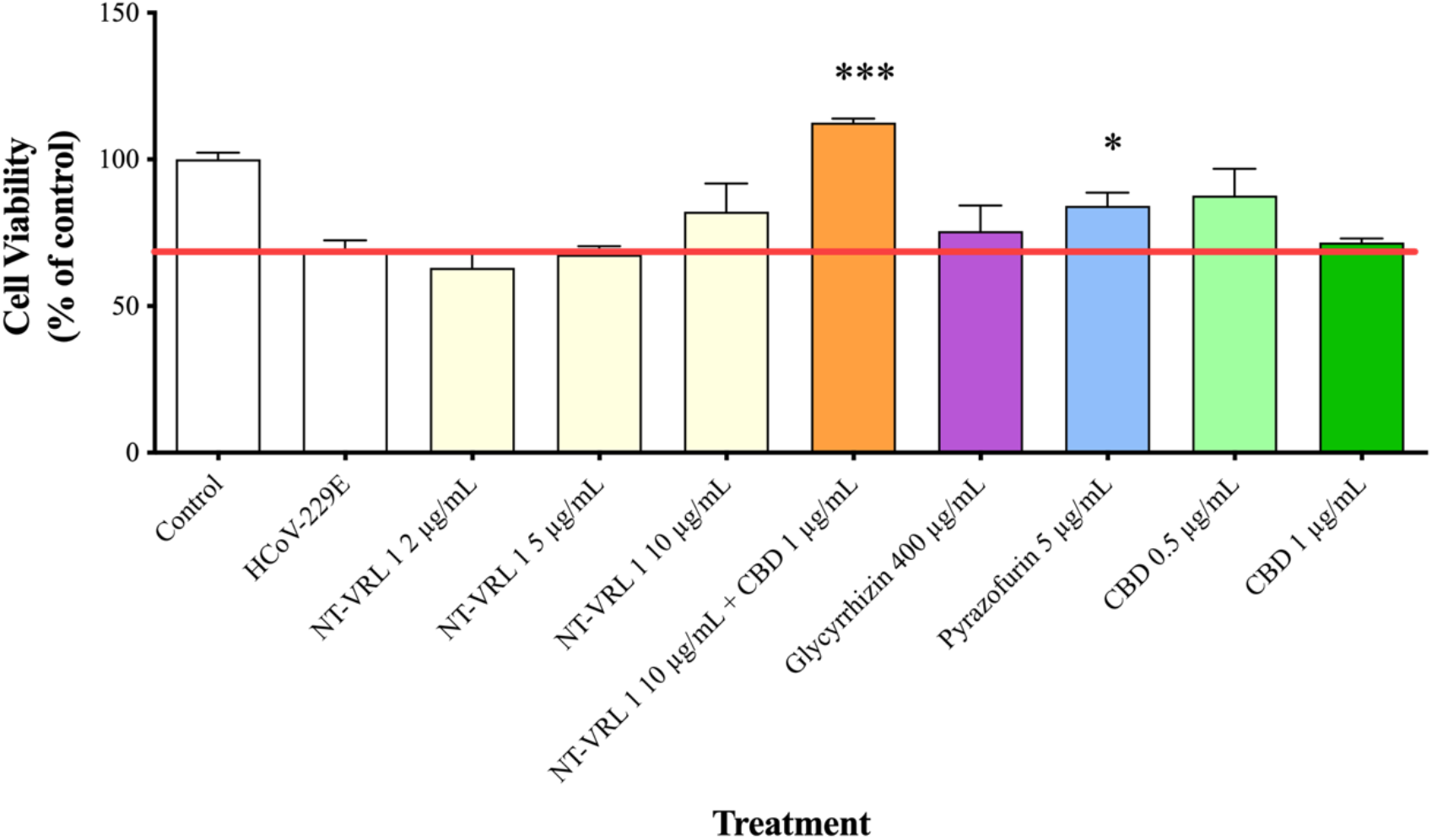
Protective effects of compounds on MRC-5 cells after virus adsorption. HCoV-229E was first added to MRC-5 cells for 1 h. Then, different concentrations of compounds were added to the cells for an additional 72 h. Cell viability was determined using an XTT assay. Results represent mean percent viability ± SEM (*n* = 4). Statistics are presented for each treatment relative to cells treated only with HCoV-229E. \**p* < 0.05 and \*\*\**p* < 0.001, according to a *t*-test.

**Figure 7:**
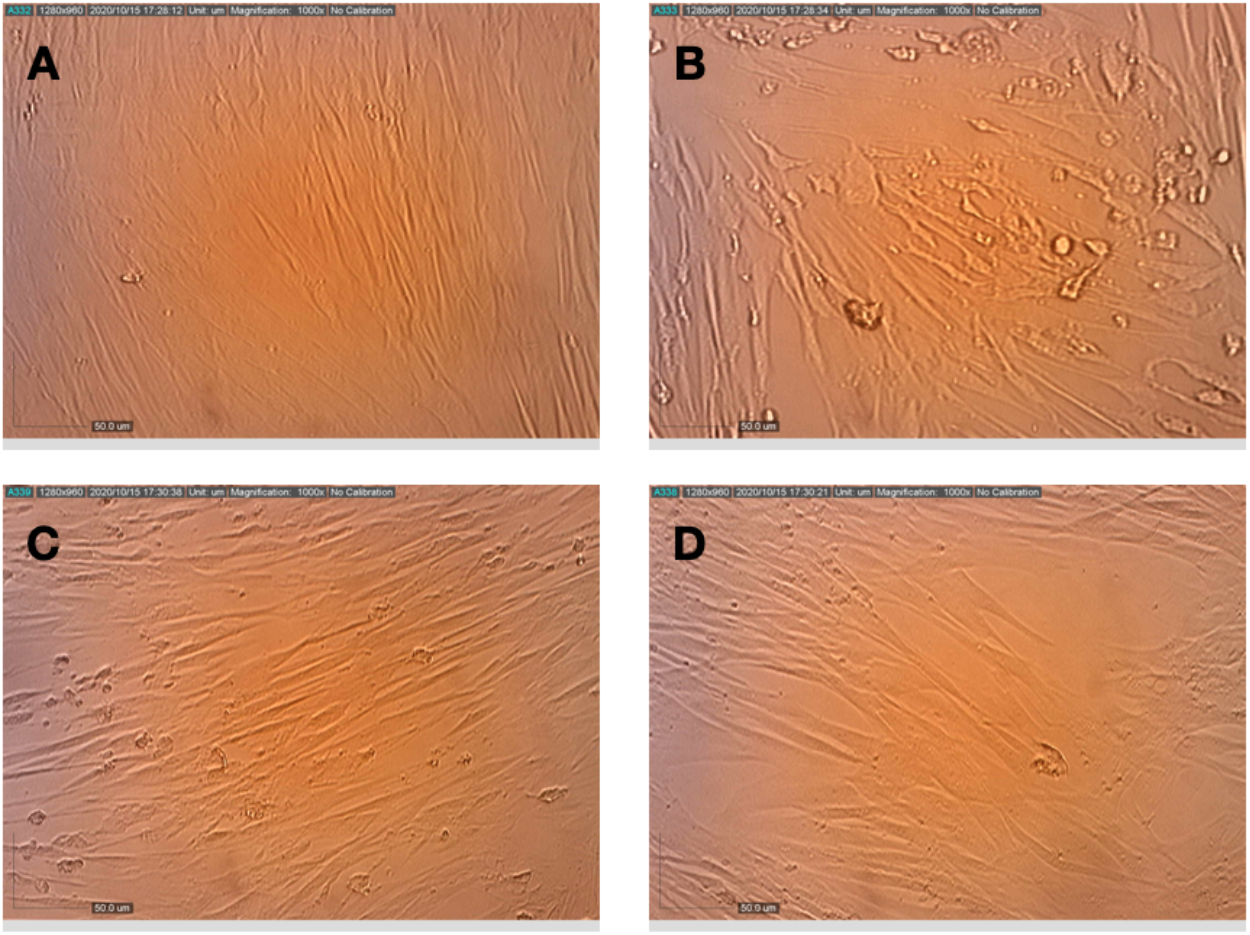
Effects of terpenes and CBD applied post-infection on the replication and cytopathic effect of HCoV-229E in MRC-5 cells. (A) Healthy MRC-5 cells, (B) MRC-5 cells at 72 h after infection with HCoV-229E, (C) MRC-5 cells that were pretreated with terpenes, photographed 72 h after infection with HCoV-229E and (D) MRC-5 cells that were pretreated with terpenes and CBD, photographed 72 h after infection with HCoV-229E.

## Discussion

Human coronaviruses have presented a great burden to global health since the 1960’s (20). The development of novel, effective antiviral solutions with low toxicity and few side effects is a matter of great interest. Secondary plant metabolites such as terpenes and cannabinoids have been shown to have significant antiviral potential and low toxicity, making them good candidates for use as antiviral agents with minimal side effects (21).

With global COVID-19 vaccination in its initial stages, the timeframe for full global vaccination is still unknown. Several population groups, such as the youngest age groups and people with health limitations, will take longer to vaccinate. Therefore, a preventative antiviral treatment to be used in conjunction with vaccines or even temporarily until vaccination or other alternatives become available would be valuable.

The objective of this study was to evaluate the anti-viral activity of the NT-VRL-1 terpene formulation, with and without CBD, against human HCoV-229E in human lung fibroblasts *in vitro*. In this study, we report the antiviral activity of the NT-VRL-1 terpene formulation and show that that activity was enhanced when it was applied together with CBD, suggesting either a synergetic or additive effect between the terpene formulation and CBD. Several studies have suggested that phytochemicals found in cannabis may be useful as potential anti-inflammatory agents (22, 23). Such activity may be particularly useful for controlling the cytokine storm syndrome and acute respiratory distress syndrome associated with COVID-19. This study is the first to test cannabis phytochemicals for use against a coronavirus.

Pyrazofurin, a natural antiviral compound that has been shown to be effective against SARS-associated coronaviruses (17), was used as positive control. Glycyrrhizin, which has been shown to have antiviral effect against SARS-associated coronaviruses (24), served as the second positive control.

The mode of antiviral action of NT-VRL-1 was determined by the addition of the compounds to uninfected lung cells, before or after those cells were inoculated with HCoV-229E. The time-of-addition assays can help us to determine the point(s) at which our compound inhibits HCoV-229E replication.

Our results demonstrate that NT-VRL-1’s antiviral effect was most pronounced in the pretreatment system, which may indicate that the compounds’ antiviral effect is based on the prevention of viral attachment and/or entry. Under these conditions, both CBD (at 0.5‒1 μg/mL) and NT-VRL-1 (at 2‒10 μg/mL) exhibited observable antiviral effects, as did the positive controls (pyrazofurin at 5 μg/mL and glycyrrhizin at 400 μg/mL). In addition, when CBD (1 μg/mL) and NT-VRL-1 (10 μg/mL) were applied together, we observed a synergistic antiviral effect that was even stronger than that observed for the positive controls.

Terpenes have been shown to have antiviral activity against SARS-CoV (17, 19). However, in previous studies, the terpenes were added to the virus at the same time and no time-of-addition assays were performed. To the best of our knowledge, this is the first report on the antiviral mode of action of terpenes and CBD against a coronavirus.

NT-VRL-1 exhibited an antiviral effect and should preferably be pre-incubated with cells prior to virus exposure. The combination of NT-VRL-1 with CBD amplified this antiviral effect. These results suggest that NT-VRL with or without CBD could be useful as a preventative measure against coronaviruses. As the lungs are the organs most affected by COVID-19, preventative treatment directly to the lungs, possibly via inhalation, would be the ideal administration route for this potential therapeutic solution.

## Acknowledgments

We thank Eybna team for their constant encouragement throughout this project. We would like to thank Ina Stelmah and Shlomit Lempert for their supportive efforts and helpful suggestions. We gratefully acknowledge the support of our work and our ideas by our partner Seach Medical Group. Finally, we thank Yaakov Amidror for the valuable discussions related to this work.

